# Uptake of aminoglycosides through outer membrane porins in *Escherichia coli*

**DOI:** 10.1101/2022.09.05.506620

**Authors:** Eshita Paul, Ishan Ghai, Daniel Hörömpöli, Heike Brötz-Oesterhelt, Mathias Winterhalter, Jayesh A Bafna

## Abstract

Aminoglycosides are important clinical antibiotics but their molecular uptake mechanism is still not completely understood. Here we quantify and compare the passive transport of three aminoglycosides (kanamycin, gentamicin, and amikacin) across general or sugar specific porins of *Escherichia coli* (OmpF, OmpC, LamB and ChiP). Our analysis revealed that permeation of aminoglycosides (Kanamycin/Gentamycin/Amikacin) is about the same through ChiP (≈5/3/2 molecules/s), OmpF (≈10/15/<1 molecules/s) and OmpC (≈11/8/<1 molecules/s). In contrast, LamB of smaller pore diameter has no significant permeation (≤1/1/1 molecules/s, all values recalculated for a gradient of 10 µM). Biological assays confirmed the relevance of these translocations for antibiotic potency.

## Introduction

Gram-negative bacteria are enclosed by the complex multi-layered cell envelope containing the cytoplasmic membrane, the peptidoglycan layer and the outer membrane ^1–3^. Among these chemically and structurally diverse layers, the outer membrane represents the first line of bacterial defense^4^. To overcome the outer membrane barrier, hydrophobic antibiotics take the slow lipid-mediated route, while the small hydrophilic ones – like β-lactams, tetracycline, chloramphenicol, and fluoroquinolones pass through general diffusion porins such as OmpF or OmpC^2^. Although most of these porins are constitutively expressed and present in open conformations allowing passive diffusion of small molecules with likely a defined exclusion limit (roughly <650 Da)^5^, there are some porins which are solute-specific. Examples of such porins in *E. coli* include, but are not limited to, maltose and maltodextrin-specific LamB^6^, sucrose specific SrcY^7^, long chain fatty acid specific FadL^8^, and nucleoside-specific Tsx^9^, along with chitooligosaccharide-specific chitoporins^10^.

Among the sugar-specific porins in *E. coli*, LamB has been extensively studied for maltosugar transport including maltodextrins as substrates. Another set of studies on sugar uptake has been performed on chitoporins (ChiP) that allow diffusion of chitooligosaccharides across the outer membrane of chitinolytic bacteria. First reported identifications of chitoporins came from marine species, including *Vibrio furnissii* (*Vf*ChiP)^10^, which uses chitin as a natural source of energy via the N-acetylglucosamine pathway, followed by the chitoporin in *Vibrio cholerae* (*Vc*ChiP)^11^. Recently, Suginta and co-workers proved the physiological function of the outer membrane chitoporin from *Vibrio harveyi* (*Vh*ChiP)^12^ by heterologous expression in *E. coli*, showing specifically a high translocation rate for chitohexose. The *chip* gene encoding chitoporin is conserved in non-chitinolytic bacterial species as well e.g. in *E. coli*^13^. Here the ChiP channel encoded by the *chiP* gene (formerly *ybfM*) has been shown to also facilitate the specific transport of chitin degradation products across the outer membrane^4^. Constitutive expression of *chiP* in *E. coli* is repressed by transcriptional and post-transcriptional control mechanisms by the ChiX sRNA^14^. In a *chiX* knock-out mutant, the presence of chitobiose/chitotriose induced the expression of *chip* resulting in increased cellular levels of both mRNA and protein ^15^. ChiP of *E. coli* shows sequence homology to the OprD family of porins and a monomeric structure is predicted ^4^. Surprisingly sequence comparison of ChiP from *E. coli* to ChiP from opportunistic pathogens like *Salmonella typhimurium*^15^ (Q7CQY4) reveals 90% sequence identity, followed by 70% identity to *Serratia marcescens* ChiP^16^ (L7ZIP1) and significantly lower identity (12-14%) to *Vibrio* ChiP sequences. At the same time ChiP of *E. coli* shows broad substrate specificity with a lower binding constant for chitooligosaccharides as compared to its *Vibrio* homologs. As per literature^4^, ChiP functions in *E. coli* only under stress conditions unlike the constitutive activity of *Vibrio* ChiPs, e.g., during the utilization of chitin as a source of energy in glucose-scarce environment. This broad specificity for substrate might further make this a potential porin for allowing the transport of molecules with sugar-like structures.

Experimental investigations on bacterial antibiotic sensitivity revealed modifications in porin profiles, including reduction in the expression level or overexpression of functionally mutated porins, as an emerging mechanism in antibiotic resistant isolates^17^. This brings the study of the antibiotic influx process into the limelight. Here we focus on aminoglycosides, which are amino sugar containing aminocyclitols, binding to 16s/30s rRNA and in turn inhibiting protein synthesis by miscoding, inhibition of translocation or inhibiting recycling of ribosomal subunits and even disrupting biogenesis of the ribosome^18–21^. Uptake of clinically relevant aminoglycosides across the cytoplasmic membrane of Gram-negative bacteria is a two-step energy dependent process and prior to that, additional transport across outer membrane has to be achieved^22–24^. These polycationic aminoglycosides have been shown to follow the species-specific self-promoted uptake mechanism^25^. The polycations displace divalent cations cross-bridging neighboring LPS molecules by competitive binding, which results in destabilization and permeabilization of the outer membrane^25^. Our recent study^26^ demonstrated the contribution and relevance of the major outer membrane porins OmpF and OmpC in the molecular uptake of kanamycin, one of the most abundantly used aminoglycosides, into *E. coli*, showing a new diffusion route of these aminoglycosides through outer membrane porins.

With the sugar-like structure of the aminoglycosides in mind, we were interested whether these aminoglycosides are recognized as a substrate for translocation through the sugar specific minor porin ChiP of *E. coli*. So far only major porins have been studied in great details for their contribution to antibiotic transport^27^. To identify the contribution of this specific minor porin, we used single channel electrophysiology experiments. In the presence of low concentrations of antibiotics, we were unable to detect the translocation events due to their fast pace. We confirmed and compared the uptake of above molecules through ChiP and other porins using bi-ionic reversal potential measurements and supported the relevance using whole-cell studies with a defined set of *E. coli* porin mutants. Our interdisciplinary study showed a robust uptake pathway for aminoglycosides through ChiP of *E. coli* shedding light on their nonspecific transport through this minor porin. This study increased our understanding of the aminoglycoside translocation into Gram-negative bacteria, which might be useful for improving their uptake.

## Materials and Methods

### Bacterial strains and chemicals

For native porin overexpression, we used *E. coli* BL21 (DE3) Omp8^28^ strain lacking the major porins of *E. coli* (OmpF, OmpC, OmpA and LamB). These cells were transformed with the respective plasmids and grown at 37°C, 200 rpm in Luria-Bertani (LB) medium (Roth, Germany) or LB agar plates supplemented with ampicillin at 100 μg/mL for plasmid maintenance. Gentamicin sulfate was obtained from Carl Roth, kanamycin sulfate, amikacin disulfate, potassium chloride salt from Sigma-Aldrich (Dorset, United Kingdom), 1,2-diphytanoyl-sn-glycero-3-phosphocholine (PC) from Avanti Polar Lipids (Alabaster, AL) and all other chemicals used were procured from AppliChem.

### Gene knockout generation

For gene deletions in *E. coli*, the method by Datsenko and Wanner was used^29^. Briefly, primers with homologous sequences corresponding to the target gene were used to amplify a kanamycin resistance cassette from the pKD13 plasmid (**Table S1**). Electrocompetent *E. coli* cells containing the pKD46 plasmid were transformed with this PCR product and homologous recombination was initiated by λ Red recombinase from the pKD46 plasmid. Successful integration of the resistance cassette was checked by growth on LB with 50 µg/ml kanamycin. Gene deletion mutants were confirmed by PCR and followed by Sanger sequencing.

### Susceptibility measurements

The minimal inhibitory concentration (MIC) was determined as described by the standardized method of CLSI^30^. Briefly, an antibiotic two-fold concentration series was prepared in microtiter plates in cation adjusted Mueller Hinton broth II (MHBII), and *E. coli* cells were added to a final inoculum of 5 × 10^5^ CFU/ml. The plates were incubated at 37°C and after 18 h of incubation, the MIC was defined as the lowest concentration completely inhibiting bacterial growth. For agar plates with a linear antibiotic concentration gradient, MHBII was supplemented with 7.5 g/L agar-agar. A first layer was poured at an inclined position into square 12 × 12 cm petri dishes. After polymerization of the agar, the second layer, containing the indicated antibiotic concentration, was added at an even position. After polymerization of this second layer, agar plates were stored over night at 4°C. On the next day, *E. coli* cells were streaked on the agar concentration gradient plates from a cell suspension in 0.9% NaCl (w/v), which was prepared from *E. coli* colonies grown on MHBII agar plates and where the optical density at 600 nm (OD_600nm_) was adjusted to 0.1. Growth was measured after 18 h of incubation at 37°C.

### Expression and purification of proteins

For the overexpression of ChiP, the gene *chiP* was cloned from genomic DNA of *E. coli* BW25113. Using the primer pair chiP-pASK5 (**Table S2**), the porin gene sequence was integrated via Gibson cloning^31^ into the plasmid pASK-IBA5 (IBA GmbH), which was beforehand digested with XbaI and XhoI. This resulted in the plasmid pASK-IBA5-chiP which was further used as the template for cloning the *chiP* gene into pET19b plasmid (Addgene). This was done using the primer pairs pET19b-chiP and chiP-pET19b with the Gibson cloning method (**Table S2**). Correct insertion was confirmed by XbaI/XhoI digestion and Sanger sequencing. *E. coli* BL21 (DE3) Omp8 was transformed with pET19b-*chiP*.

For protein expression, an overnight preculture of *E. coli* BL21 (DE3) Omp8 containing pET19b-*Ec*ChiP was inoculated into Luria-Bertani medium containing 100 μg/mL ampicillin at OD_600_=0.1 and grown at 37°C and 200 rpm in Erlenmeyer flasks till optical density of the culture reached 0.6. At this point, the expression of the protein was induced with 0.5 mM isopropyl β-D-1-thiogalactopyranoside (IPTG – Sigma) and the flask was placed at 37°C, 200 rpm. After 6 hours, cells were centrifuged at 3220xg (Eppendorf centrifuge 5810 R, A-4-62 rotor) for 30min at 4°C. The pellet was stored at -20°C for the extraction steps. For extraction, a pellet from 500 ml of culture was resuspended in 10 ml lysis buffer (20 mM Tris pH 8, 2.5 mM MgCl_2_, 0.1mM CaCl_2_, 1 mM PMSF, 10 µg/ml RNase A and 10 µg/ml DNase I) and disrupted by five passages at 15000 psi in a French Press on ice. After removing the cell debris by 30min of centrifugation at 3220xg, 4°C, the membrane was pelleted by ultracentrifugation (1 hour, 100000xg at 4°C). The pellet was solubilized in 20 mM Tris/HCl pH 8, 0.15 % Octyl POE (N-octylpolyoxyethylene from Bachem) using a Potter homogenizer and incubated for 1 hour at 4°C on the wheel. The remaining insoluble material was separated by a second ultracentrifugation (100000xg, 4°C, 1 hour). ChiP was extracted by solubilization of this pellet with 20 mM Tris HCl pH 8, 3% Octyl POE and elimination of insoluble material by ultracentrifugation. This step was performed two times. The last supernatants containing the expected protein were pooled and concentrated with an Amicon concentration unit (cut off 30 kDa). The samples were supplemented with Laemmli sample buffer and heated at 95°C for 10 min (denaturing condition) or supplemented with Laemmli sample buffer lacking beta mercapto-ethanol (non-denaturing condition) and loaded on 12% SDS PAGE gel. The gels were stained with Coomassie Brilliant Blue G250. To obtain pure protein from the last supernatant, the concentrated sample was subjected to anion-exchange chromatography using a Mono Q® 5/50 GL prepacked column (5.7×1ml) connected to BioLogic DuoFlow chromatography system (Bio Rad). Proteins were eluted with a linear gradient of 0-1 M KCl in 20mM phosphate buffer, pH 7.4 containing 0.2% (v/v) lauryldimethylamine N-oxide (LDAO) as previously reported^4^. The purity of the ChiP was confirmed by SDS-PAGE analysis and on our gels the native protein showed a slightly higher molecular mass than in the former study^4^. The presence of the protein was confirmed by mass spectrometric analysis and Western blot analysis using an anti-ChiP antibody. OmpF and OmpC were expressed and purified according to the previously published protocol^32,33^.

### Planar lipid bilayer and electrical recording: Single and multichannel measurements

Planar lipid bilayers conferring to Montal and Mueller were formed as published^26,34^. Briefly, an aperture in a Teflon septum with a diameter of 100-120 μm was pre-painted with hexadecane dissolved in n-hexane at 1-5% (v/v) and the cuvette compartments were dried for 10-15 min, in-order to eliminate the solvent. Bilayers were made with 1,2-diphytanoyl-sn-glycero-phosphatidyl-choline at a concentration of 4-5 mg/ml in n-pentane. Stock solutions (0.5 mg/mL) of the outer membrane porins were diluted 10^3^ to 10^4^-fold using Genapol X-080 (1% v/v). A small volume from the dilution was added to the *cis* side of the chamber containing 2.5 ml of 1M KCl for single channel measurements. For reversal potential measurements the gradient was created with the pure antibiotic solution^26,36^. Throughout the experiments the pH was maintained at 7.0 ± 0.5. The *cis* side of the chamber was the virtual ground and the *trans* side was linked with a CV-203BU Headstage of the Axopatch 200B (Molecular Devices, LLC) patch-clamp amplifier in V-clamp mode (whole cell β = 1). The output signal was filtered by a lowpass Bessel filter at 10 kHz and saved at a sampling frequency of 50 kHz using an Axon Digidata 1440A digitizer (Molecular Devices, LLC). Data analysis was performed with an in-house analysis suite created with the LabVIEW software suite (National Instruments) or using Clampfit (Axon Instruments). Standard Ag/AgCl electrodes were used to detect the ionic current in single channel experiments. For measuring Bi-ionic and tri-ionic reversal potentials (multichannel) to allow for asymmetric condition, we used commercial calomel electrodes (Metrohm) containing a salt bridge. The current voltage relation of the individual experiments was calculated from single averaged currents at the given voltage. The relative permeability of cations *vs* solute anions in the bi-ionic case (P_antibiotic_^+^/P_sulfate_^−^) and in tri-ionic case (P_K_^+^/P_sulfate_^−^/P_antibiotic_^+^) were obtained by fitting of the experimental I−V curves with the Goldman−Hodgkin−Katz^35^ current equation^36^.

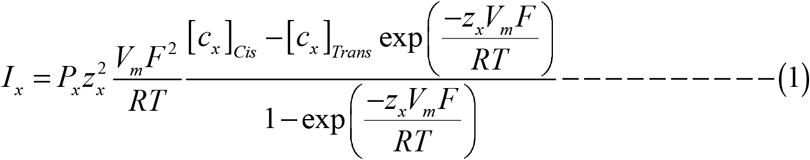

## Results

### Single channel and bi-ionic reversal potential measurement

In a series of measurements, we took the single channel conductance of OmpF, OmpC, ChiP and LamB in Kanamycin 20mM, Gentamicin 20mM and Amikacin 5mM respectively. Moreover, antibiotic ions have much lower conductances and thus the single channel conductance values for LamB are only estimates (**Table 1**). Further the single channel conductance of OmpF, OmpC, ChiP, LamB were also measured for 1M KCl (**Table S3**) which are in agreement with previous measurements^6,26,32,33^. For this we prepared lipid membranes in the respective electrolyte solution and added the respective protein on one side, usually the ground connected side. The subsequent increase in conductance steps were statistically analyzed and the smallest conductance peak was identified as single channel conductance.

**Table 1.** Measured single channel conductance in presence of antibiotic salt under symmetric condition and reversal potential measurements for the indicated concentration gradient (5 and 20 mM). From these values we extrapolated the permeability ratio and the flux. Details of the flux calculations are outlined in the supplement.

| Concentration gradient of Kanamycin $\Delta = 20$ mM | | | | |
| --- | --- | --- | --- | --- |
| & $V_{rev}$ | | Conductance estimated | $P = P_{cation}^{4+} / P_{sulfate}^{2-}$ | (molecules/sec/monomer)<br>(Extrapolated to 10 μM gradient) |
| OmpF | | 16 ± 5 pS 9.6 mV | 1:14 | 10 |
| OmpC | | 13 ± 3 pS 5.1 mv | 1:4.2 | 11 |
| Chip | | 7 ± 3 pS 6.5 mV | 1:6 | 5 |
| LamB | | 3 ± 2 pS not measured | $\ll 10^7$ | $\leq 1$ |
| Gentamycin $\Delta = 20$ mM | | | | |
| | | Conductance estimated | $P = P_{cation}^{4+} / P_{sulfate}^{2-}$ | |
| OmpF | | 24 ± 8 pS 15 mV | 1:25 | 15 |
| OmpC | | 15 ± 5 pS 16.5 mV | 1:30 | 8 |
| Chip | | 10 ± 4 pS 2 mV | 1:3 | 3 |
| LamB | | 4 ± 1 pS not measured | $\ll 10^7$ | $\leq 1$ |
| Amikacin $\Delta = 5$ mM | | | | |
| | | Conductance estimated | $P = P_{cation}^{4+} / P_{sulfate}^{2-}$ | |
| OmpF | | 14 ± 9 pS | 1:10<sup>6</sup> | $\leq 1$ |
| OmpC | | 9 ± 5 pS | 1:10<sup>6</sup> | $\leq 1$ |
| Chip | | 5 ± 4 pS 10 mV | 1:10 | 20 |
| LamB | | 2 ± 1 pS not measured | $\ll 10^7$ | $\leq 1$ |

In a second series of measurements, we made the lipid membrane in 1M KCl and inserted the respective single channel. Addition of small amounts of antibiotics resulted in channel blockages. A statistical analysis of the blockage lengths or dwell times as a function of external voltage is used to reveal an estimate for possible translocations through the channel. To begin with, the three porins of interest from the *E. coli* outer membrane (OmpF, OmpC, and ChiP) were reconstituted into a planar DPhPC bilayer (in individual experiments) from the electrically grounded or the *cis* side, and the single channel currents in the presence of 1M KCl at pH 7 were recorded. Single channel blockages were observed at a concentration of 100 μM gentamicin for ChiP when added to the *cis* side of the chamber, and a transmembrane potential of -100mV was applied on the *trans* side. (**Figure 1**). Such behavior is due to the cationic nature of gentamicin and with the application of negative potential, the molecules are pulled through the pore due to the electrophoretic force. Decreasing the magnitude of ΔV further decreased the number of blockages, as the gentamicin molecules were pulled at a slower rate across the channel by virtue of a weaker electrophoretic force. To quantify the findings, the event rate f_e_ was analyzed, which is the number of events per second, and the dwell (or residence) time of gentamicin molecules in the channel was analyzed as obtained from an exponential fit of the dwell time distribution (**Figure 2**). For both quantities, multiple events were analyzed for each transmembrane voltage applied in the range from -20 to -100 mV. In a similar manner, measurements were also performed for OmpF and OmpC, where adding 10 μM of gentamicin was enough to observe a significant number of blockages (**Figure 1**). Again, the event rates and dwell times were analyzed for these two porins, but we did not observe any trend in dwell time change, although raising the negative voltage increased the overall number of events. As mentioned in our previous paper, an event rate itself is not sufficient to conclude translocation and a decrease of the dwell time with higher voltage is a hallmark of translocation^26^. The independent behavior of dwell time in the given range of transmembrane voltage indicated for OmpF and OmpC, that most events are rather simple blockages and not actual translocations.

**Figure 1.**
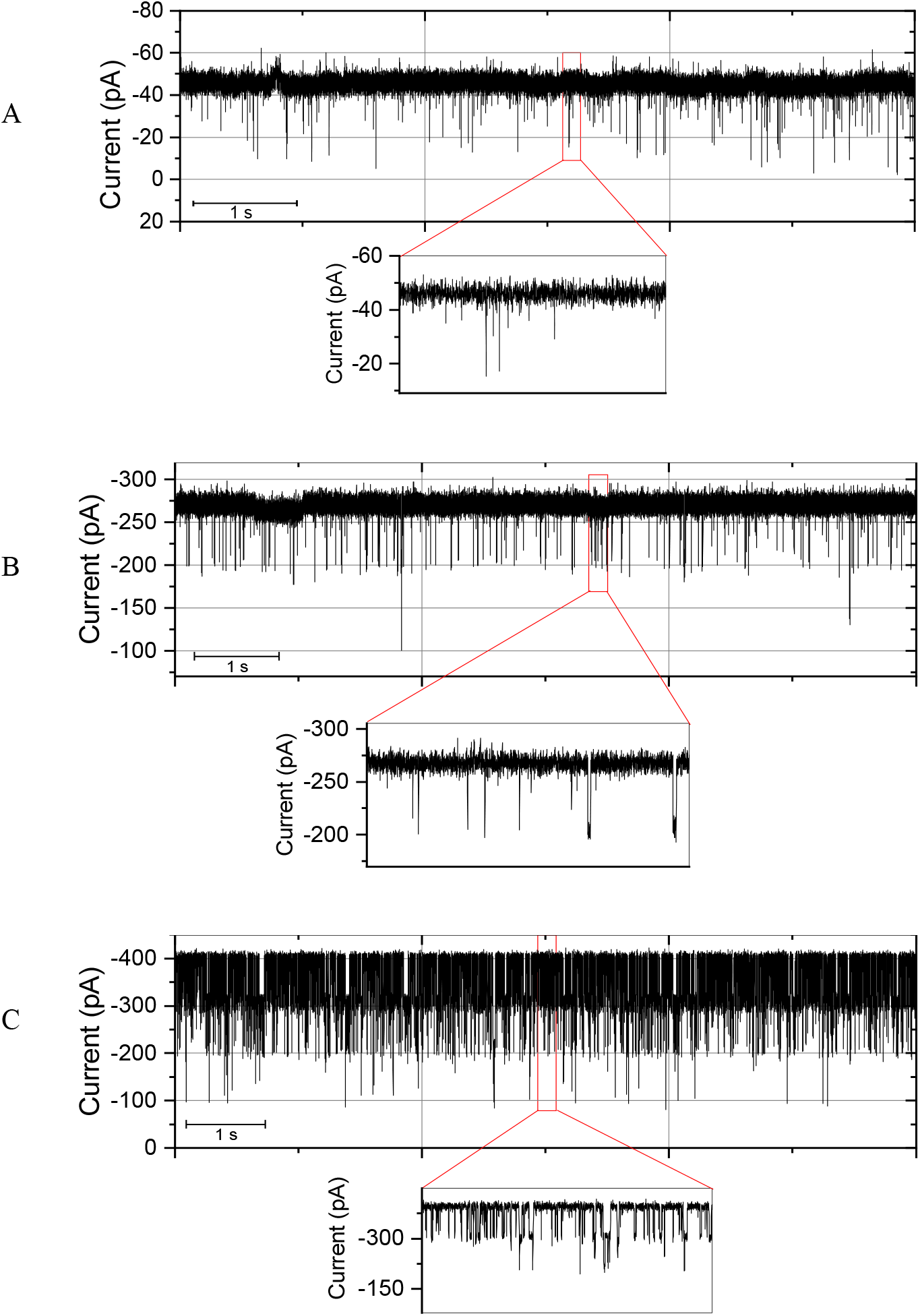
Examples of ion current traces through a single porin channel in the presence of gentamicin at - 100 mV applied transmembrane volatge. **A:** ChiP in presence of 100 μM, **B:** OmpC in presence of 10 μM, and **C:** OmpF in the presence of 10 μM gentamicin sulfate in 1 M KCl, 10 mM HEPES at pH 7. The insets show zoomed events. All experiments were performed at 25°C while the data were filtered at 1 KHz.

**Figure 2.**
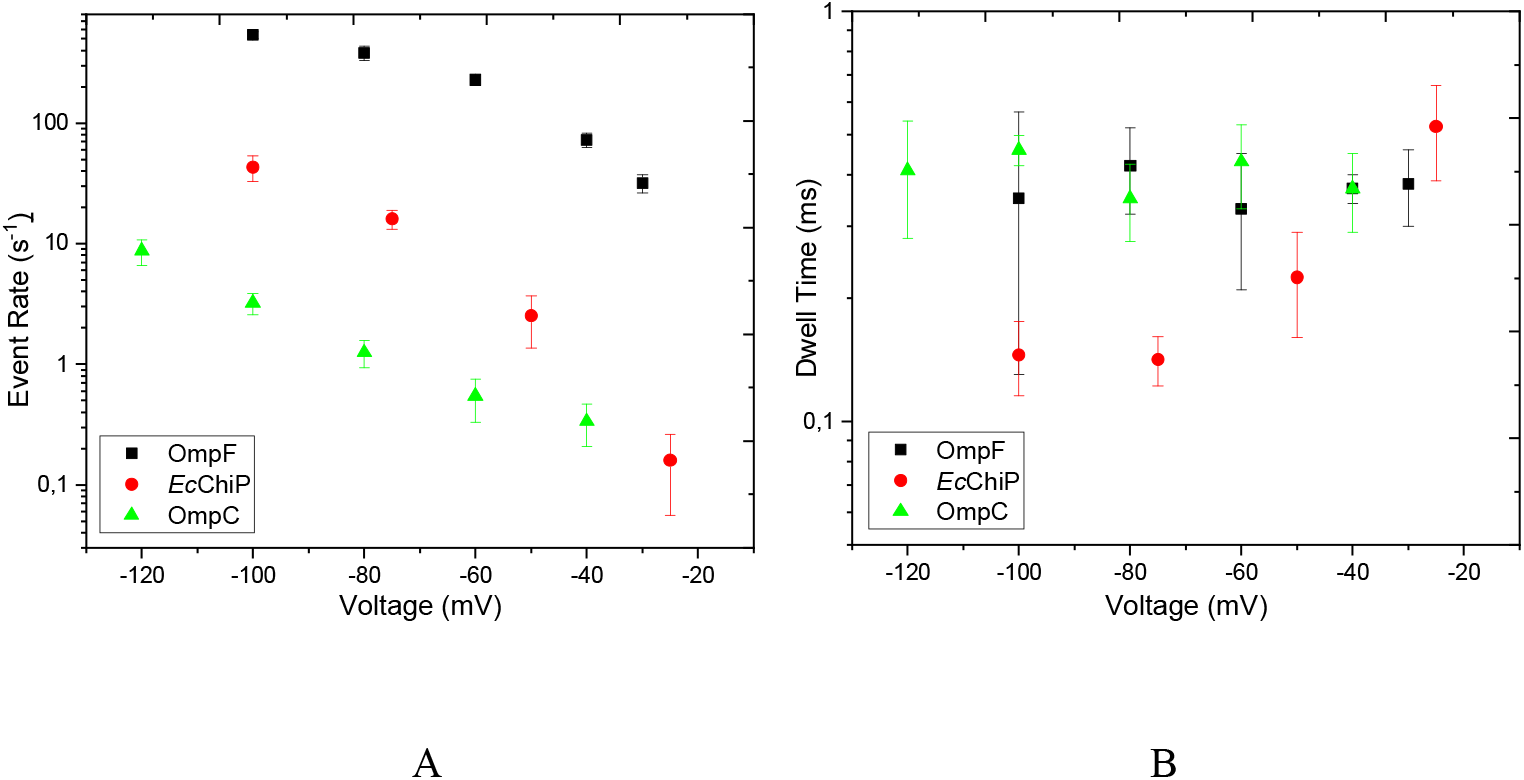
Event rates and dwell times for gentamicin. Statistical analysis of the traces measured at different applied voltages for OmpF, OmpC, and ChiP. **A:** Event rate f_e_ *vs* applied negative voltages. Increasing the negative voltage pulls more cationic molecules into the channel. **B:** Dwell time analysis. In the case of ChiP, higher negative voltage drives gentamicin faster through the channel. In contrast, for OmpF and OmpC, increasing the voltage in the range of -20 to -100 mV does not affect the residence time significantly and the dwell time is independent of the voltage, indicating mainly non-translocating events or an overestimation of possible translocation.

In a third series of measurements, we followed a different approach previously used to reveal translocation for charged compounds as the current fluctuation analysis does not always identify successful transport by distinguishing translocation from binding or reflected events. To directly characterize quantitative transport of such charged molecules, the zero current assay or bi-ionic reversal potential assay is used. To distinguish permeability from short-lived bounce back events, we performed a concentration-driven relative permeation assay using bi-ionic reversal potentials for all the three antibiotics for each of the porins (**Table 1, Figure S2**). We produced a symmetrical bi-ionic antibiotic-sulfate condition on both sides of the porin reconstituted bilayer. A concentration gradient was then established by increasing the solute concentration on the *cis* (ground) side of the bilayer, and the induced zero-current membrane potential was observed. The permeability ratio of the antibiotic: sulfate through the OmpF, OmpC and ChiP were determined using the simplified GHK equation considering the effective charge of an antibiotic molecule as +4 for this bi-ionic situation. In this case, the reversal potentials for all the antibiotics are negative for the same ionic concentrations cis/trans for all the three porins, implying that anions permeate faster compared to cations through the three porin channels OmpF, OmpC and ChiP (**Table S5**).

For gentamicin the permeability ratios obtained were 1:25, 1:30 and 1:3 for OmpF, OmpC and ChiP. These ratios imply that while both sulfate and gentamicin molecules can permeate through these channels, for each gentamicin cation, the sulfates anions are permeating roughly 25, 30 and 3 times faster, respectively, until this flux is balanced by the reversal potential. This reversal potential approach is relatively simple to measure and provides a handle to qualitatively compare the translocation rate of a specific analyte across different porins. Further calculations from these measurements indicate that flux for gentamycin molecules is in a same range for ChiP as compared to OmpF and OmpC. While for kanamycin the flux rates are higher for ChiP as compared to OmpF and OmpC, whereas for amikacin, the rate of translocation is significantly higher through ChiP than through the other porins. In this context, as a negative control we also checked for the permeation of these aminoglycosides through the major sugar specific porin of *E. coli* – LamB. The reversal potential value of V_rev_ within experimental error, measured in a tri-ionic conditions clearly shows that these large molecules cannot permeate through LamB. In **Table S4** supplementary file, calculated permeability ratios obtained under tri-ionic conditions are listed for kanamycin sulfate, gentamicin sulfate and amikacin sulfate respectively.

### Whole cell assay

To validate the biological relevance of our *in vitro* experiments, we tested the antibiotic susceptibility of different *E. coli* porin mutants that had been constructed in the background of *E. coli* BW25113, a K-12 derived, broadly-used laboratory model strain. To quantify the antibacterial activity of the aminoglycosides, we measured the minimal inhibitory concentration (MIC) by a guideline-conform standardized method in liquid culture in microtiter plates (**Figure 3 A, B**, and **C**). The MIC results were complemented by streaking the *E. coli* strains on an agar plate containing a linear antibiotic gradient and measuring their distance of growth (**Figure 3D, E, F**). The deletion of only one of the major porin genes, i.e., either *ompF* or *ompC*, did not increase resistance of *E. coli* BW25113 against kanamycin, amikacin or gentamicin in the gradient agar assay, and the differences in the MIC assay were also weak. However, the combined deletion of both *ompF* and *ompC* led to a significant reduction of susceptibility in the gradient agar assay. In the case of *chiP*, already the single gene deletion increased resistance against all three aminoglycosides tested, and the effects were clearly visible on agar concentration gradient plates as well as in the MIC assay, which supports the involvement of the ChiP channel in translocation of this compound class. Notably, the overall impact of the *chiP* single gene deletion was even stronger than that of the combined Δ*ompF*Δ*ompC* double gene deletion, suggesting that ChiP plays a physiological role in the passage of all three aminoglycosides into *E. coli* BW25113. The *chiP* gene deletion was also tested within the *E. coli* ATCC 25922 strain background, which is a clinical isolate but, interestingly, in this strain the gene deletion had no effect on the resistance against kanamycin, amikacin or gentamicin when compared to its isogenic wild type strain (**Figure S3**). Differences in general outer membrane composition, the particular lipopolysaccharide structure, which differs between K12 strains and clinical isolates, or *chiP* expression might explain why different *E. coli* strain backgrounds are affected differently by the *chiP* gene deletion^42^.

**Figure 3.**
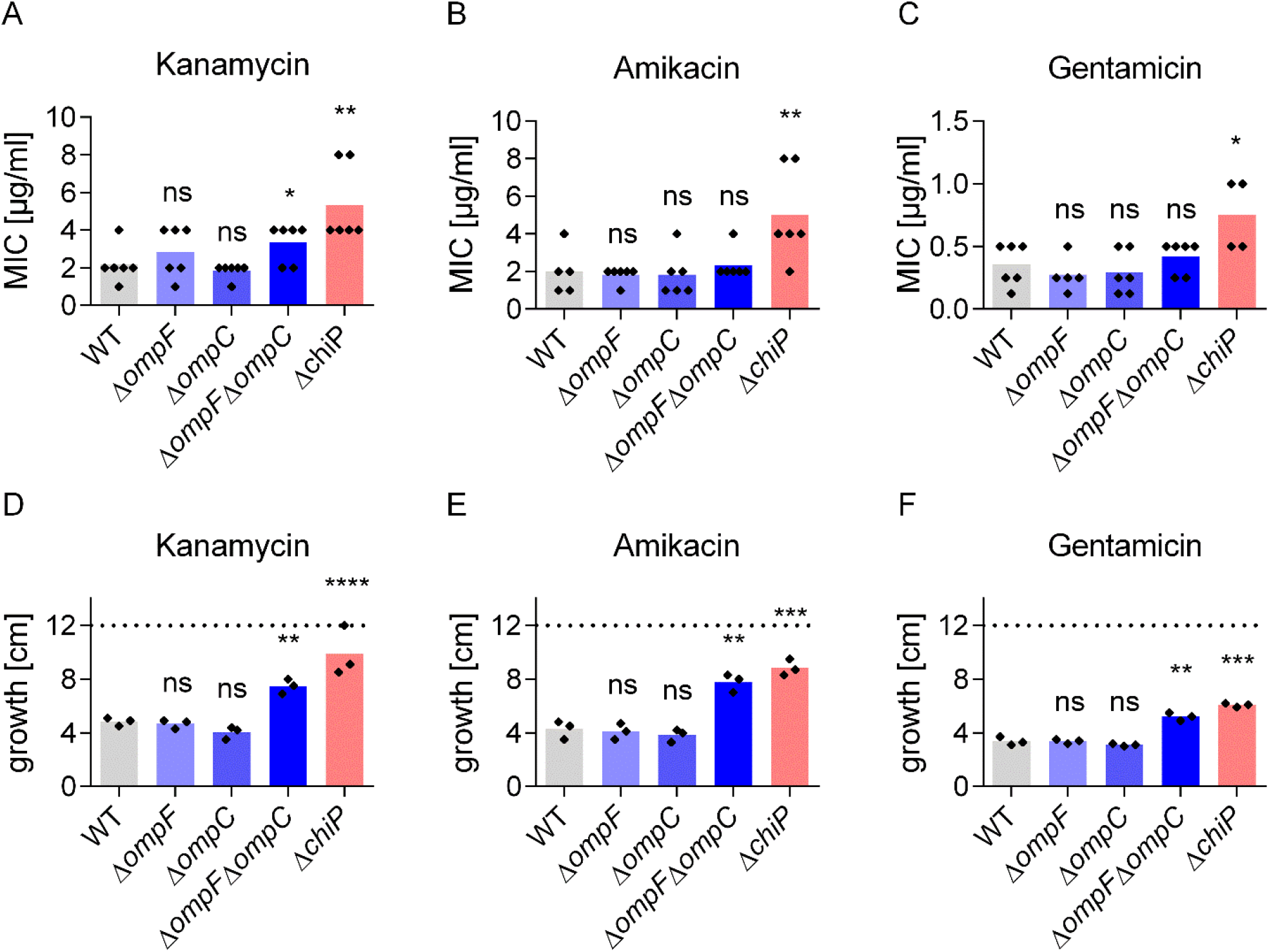
Susceptibility of *E. coli* BW25113 against the aminoglycosides kanamycin, amikacin or gentamicin. Aminoglycoside activity was measured by MIC determination (**A, B & C**) or growth on agar plates with a linear concentration gradient from 0 µg/ml to 3 µg/ml kanamycin (**D**), 0 to 2 µg/ml amikacin (**E**) or 0 to 0.75 µg/ml gentamicin (**F**). The limit of detection (i.e., the maximal stretch that a strain could grow given bei the breadth of the agar plate) is marked with a dotted line. Each diamond represents an independent experiment and the mean average is shown as a colored bar. Statistical analysis was done using unpaired Student’s t-test with Holm-Bonferroni correction, comparing the mutants to the wildtype (WT) strain. Not significant (ns), P > 0.05; **, P ≤ 0.01; ***, P ≤ 0.001.

## Discussion

The aim of this study was to quantify the possible flux of aminoglycosides through a few main *E. coli* porins. The electrophysiology technique provided quantitative information on the rate of transport and the nature of interaction at a single channel level. Furthermore, we combined *in vitro* techniques with bacterial growth inhibition assays to compare the permeation of three selected aminoglycosides side-by-side across several membrane channels located in the outer cell membrane of *E. coli*. Our data indicates that kanamycin, amikacin and gentamicin can all translocate through ChiP of *E. coli*. The permeation of kanamycin through OmpF and OmpC was already previously characterized^26^ and revisited here. According to the reversal potential data presented here, OmpF and OmpC are more restrictive to gentamicin compared to ChiP and amikacin is even more strongly preferred by ChiP in relation to the other two porins. This special capacity of ChiP to allow the passage of aminoglycosides was also reflected in our cell-based assays. Among the three porins investigated in this study, only ChiP caused a consistent and clearly significant resistance phenotype upon single gene deletion in all of our assays. In the absence of the structural data, we can conclude that due to the sugar-like structure of these aminoglycosides, they are selected as a nonspecific substrate for the Chitoporin, which has not been shown to permeate any antibiotic previously. Although LamB represents the major sugar-specific porin in *E. coli*, it is poorly permeable to aminoglycosides. Thus, it is interesting that the permeation of these antibiotics is prominent through the minor porin ChiP as compared to the major ones, and even more so as antibiotic susceptibility loss was reproducibly associated with *chiP* single gene deletion in *E. coli* BW 25113. The reason, why this physiological phenotype only emerged in the K12 derivative *E. coli* BW 25113 is currently elusive and requires further extensive study. If the substantially shorter O-specific side chains inherent to the laboratory K12 linage plays a role is currently not known ^42^.

In this context it is also important to note that aminoglycosides have also porin-independent means of entering *E. coli*. A commonly assumed way for aminoglycoside penetration is their lipid mediated self-promoted uptake by destabilizing the outer membrane. Multiple options for cell entry may also explain, why aminoglycoside resistance is commonly associated with aminoglycoside modifying enzymes or 16S rRNA methyltransferases but not uptake mutations^37^.

Nevertheless, inspection of the literature shows that the aminoglycoside class of antibiotics can also be affected by the porin mediated resistance^37^. For other antibiotic classed (carbapenems, cephalosporins multiple porins including OmpF, OmpC have been shown to play a role in developing resistance although they seem functionally redundant.^38–40^ Most of the available studies are about the major porins OmpF or OmpC and only few reports on minor porins (e.g. NmpC) or specific porins (LamB) are available^41^. Our report on ChiP emphasizes that role of specific and minor porins concerning resistance development.

Here, we were able to demonstrate that at least three different aminoglycosides utilize the *E. coli* porin ChiP, which has not previously been linked to outer membrane antibiotic translocation. So far, the only known physiological role of ChiP in *E. coli* is chitooligosaccharide uptake under glucose-deficient conditions and ChiP was reported to be expressed at a low level in the absence of any inducer^4^. Although it is not as abundant as OmpF or OmpC, ChiP seems to contribute to aminoglycoside uptake, potentially due to its fast nonspecific permeation of the aminoglycosides investigated here. Especially considering the miscoding activity of the aminoglycosides and the common model of their entry in two phases^24^, a minor porin with high permeation capacity could make a significant contribution. That means, a comparably small number of molecules entering first may be sufficient to trigger the production of non-functional membrane proteins by miscoding, thereby destabilizing the cell envelope to compound entry in a more general way. Present whole cell data also showed a difference in the effect of porin deletions between a nonpathogenic model strain and a clinical strain of *E. coli*, which emphasizes the need for follow-up studies on aminoglycoside uptake pathways in pathogenic isolates.

## Acknowledgements

The Authors would like to acknowledge Dr. Claudio Piselli for experimental help with pET19b-*Ec*Chip expression plasmid generation. We received support from the JPIAMR network RESET-ME (BMBF-ERANET JPIAMR - 01KI1827B) and the JPIAMR Virtual Institute Translocation-Transfer 01KI1828. Moreover, financial support by the German Federal Ministry of Education and Reseach, though project Gram-Neg-Design, and by the German Center for Infection Research is gratefully acknowledged.

## Author Contributions

E.P. and M.W., D.H. and H. B.-O. conceptualised and designed experiments. E.P prepared the proteins and performed single channel experiments. J.A.B performed and analysed bi-ionic reversal potentials and conductance. I.G. performed and analysed tri-ionic reversal potentials. D.H. performed and analysed whole cell assays. E.P. and M.W. wrote the article and I.G. J.A.B. D.H. and H.B.-O. edited the manuscript.

