## Supplementary material for "Uptake of aminoglycosides through outer membrane porins in *Escherichia coli*": SI

**Figure S1.**  Ion current traces in the absence and presence of gentamicin. Single channel measurements of ChiP in the A absence or B presence of 100 μM gentamicin sulfate in 1 M KCl, 10 mM HEPES at pH 7± 0.5 at an applied potential of −100 mV.

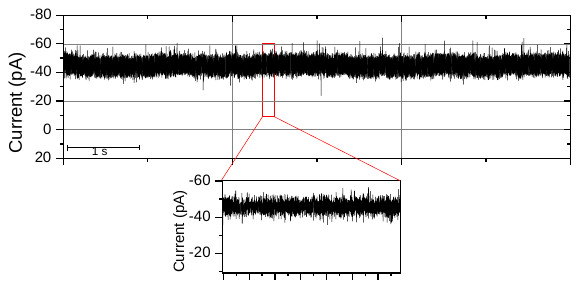

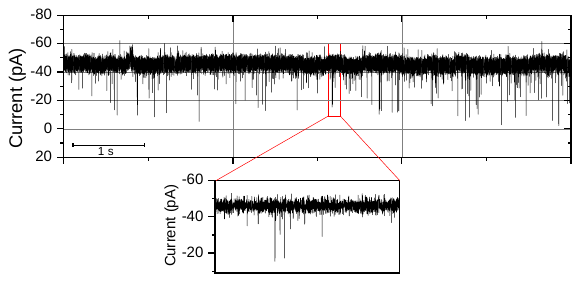

A

B

**Figure S2.** Selected I–V curves from bilayers containing multiple reconstituted ChiP channels with different aminoglycosides under bi-ionic conditions mentioned in **Table 1**. A: kanamycin **B:** gentamicin **C:** amikacin.

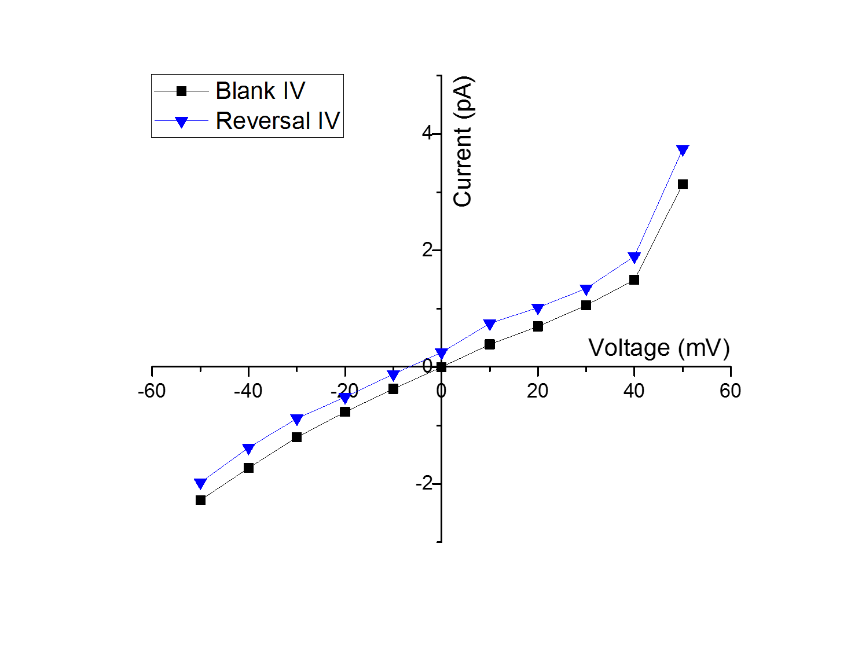

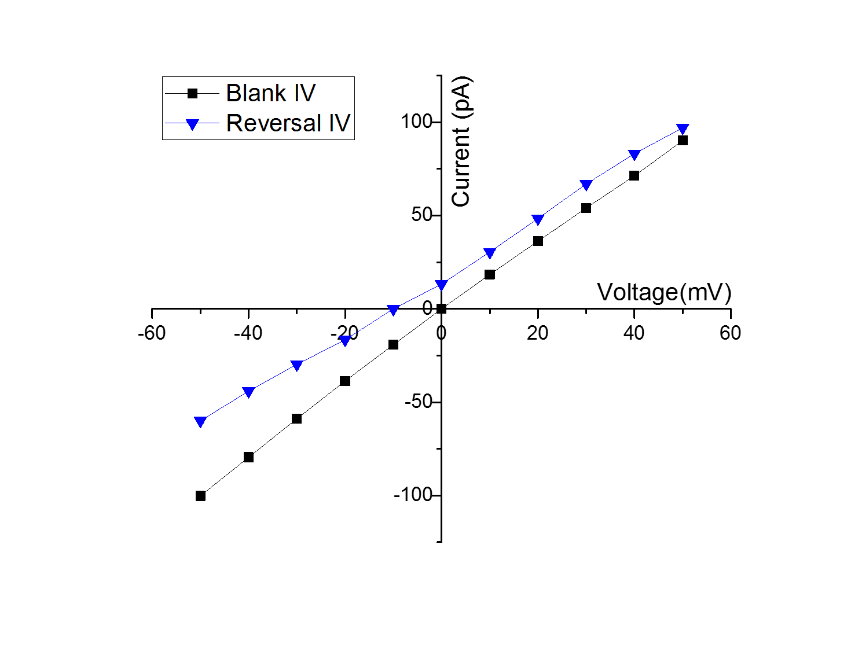

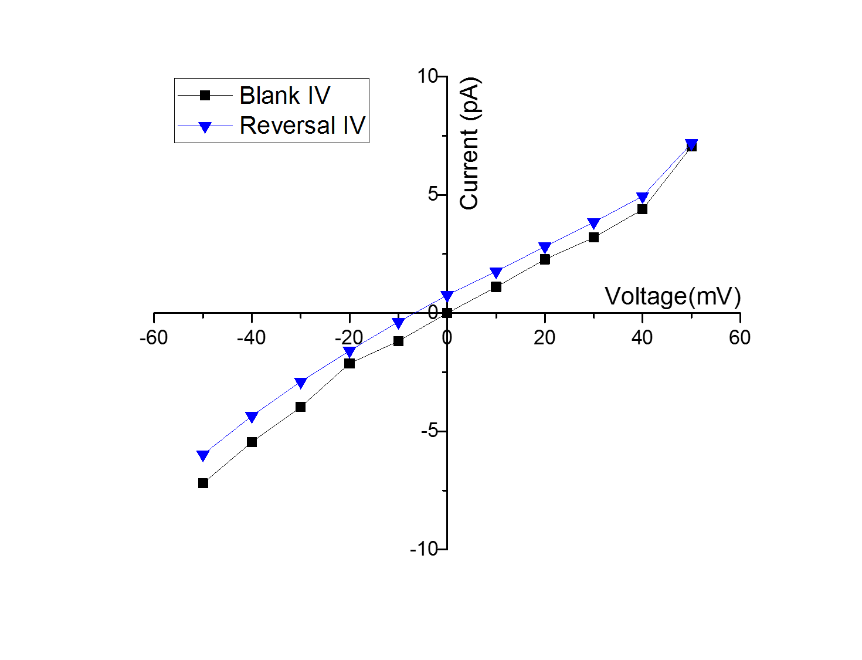

A

B

C

**Figure S2** (continued). Selected I–V curves from bilayers containing multiple reconstituted OmpF/OmpC channels with different aminoglycosides under the bi-ionic conditions mentioned in **Table 1.** **D:** OmpF and gentamicin **E:** OmpF and amikacin **F:** OmpC and gentamicin **G:** OmpC and amikacin.

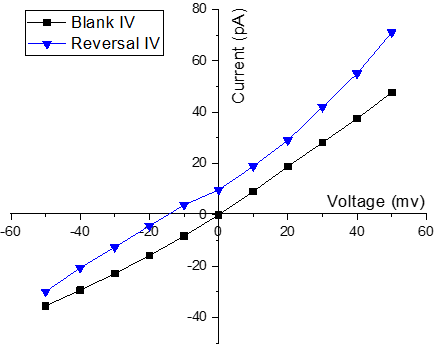

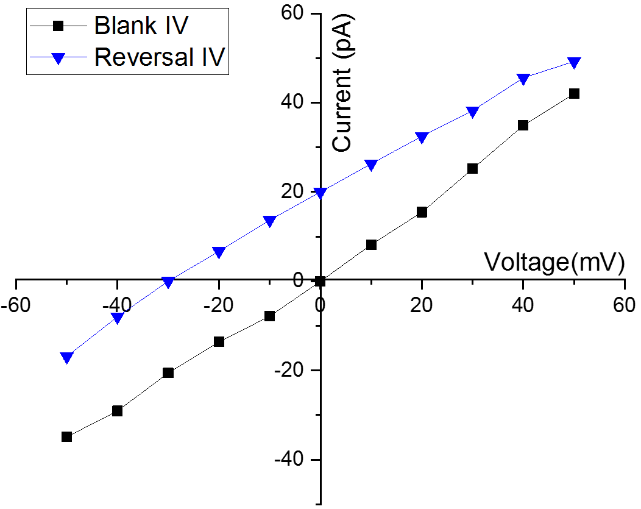

D

E

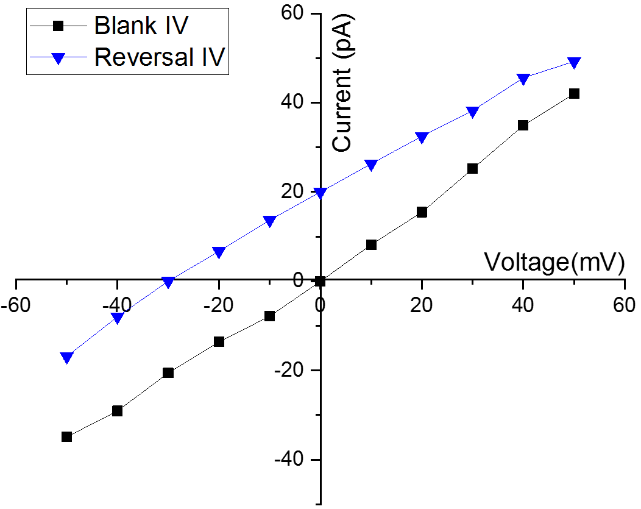

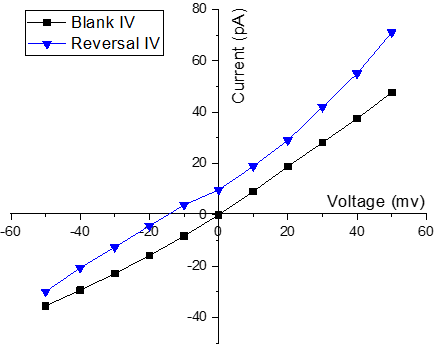

F

G

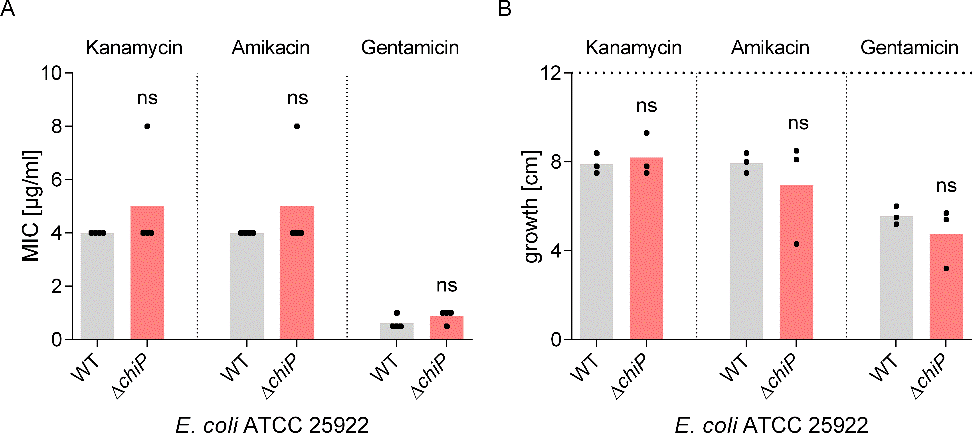

**Figure S3.** Susceptibility of *E. coli* ATCC 25922 against kanamycin, amikacin or gentamicin. A, In this strain background, the MIC determinations showed no difference between the wild type and *chiP* gene deletion strain for all aminoglycosides tested. B, *E. coli* ATCC 25922 and its isogenic Δ*chiP* mutant were streaked on an agar gradient plate with a linear concentration from 0 to 3 µg/ml kanamycin, 0 to 2 µg/ml amikacin or 0 to 0.75 µg/ml gentamicin. The gene deletion of *chiP* did not lead to growth of *E. coli* ATCC 25922 at higher antibiotic concentrations. Independent measurements are displayed as diamonds and the mean average is shown as a bar. The dotted line represents the limit of detection of the agar plates.

**Table S1.** Bacterial strains used in this study. *E. coli* BW25113 single deletion mutant strains are originated from the Keio collection^2^. For multiple gene deletion strains, the kanamycin resistance cassette was removed.^1^

| strain | genotype/characteristics | reference |
| --- | --- | --- |
| *E. coli* BW25113 | wildtype; F-, Δ(*araD-araB*)*567*, Δ*lacZ4787*(::*rrnB*-3), *λ*-, *rph*-1, Δ(*rhaD-rhaB*)*568*, *hsdR514* | ^1^ |
| *E. coli* BW25113 Δ*ompF* | F-, *Δ(araD-araB)567*, *ΔlacZ4787*(::*rrnB*-3), *λ^-^*, *ΔompF746*, *rph-1*, *Δ(rhaD-rhaB)568*, *hsdR514* | ^1^ |
| *E. coli* BW25113 Δ*ompC* | F-, *Δ(araD-araB)567*, *ΔlacZ4787*(::*rrnB*-3), *λ^-^*, Δ*ompC*768, *rph-1*, *Δ(rhaD-rhaB)568*, *hsdR514* | ^1^ |
| *E. coli* BW25113 Δ*chiP* | F-, Δ(*araD-araB*)567, Δ*lacZ*4787(::*rrnB*-3), λ-, Δ*chiP*729, *rph-1*, Δ(*rhaD-rhaB*)568, *hsdR*514 | ^1^ |
| *E.coli* BW25113 Δ*ompF*Δ*ompC* | F-, *Δ(araD-araB)567*, *ΔlacZ4787*(::*rrnB*-3), *λ^-^*, *ΔompF746*, ΔompC768, *rph-1*, *Δ(rhaD-rhaB)568*, *hsdR514* | ^3^ |
| *E. coli* ATCC 25922 | Clinical isolate, CLSI reference strain | ^4^ |
| *E. coli* ATCC 25922 Δ*chiP* | Δ*chiP* |  |

**Table S2.** Primers used for knockout and expression strain generation in this study. Nucleotides marked in bold are homologous sequences flanking the kanamycin resistance cassette of plasmid pKD13^1^.

| Oligonucleotide | Intended purpose | Sequence (5‘-3‘) |
| --- | --- | --- |
| Knockout generation | | |
| chiP_for | Deletion of *chiP* in *E. coli* BW25113 and *E. coli* ATCC 25922 | TTGGTGCAGCAATTTATACGTCAAAGAGGATTAACCCATG**ATTCCGGGGATCCGTCGACC** |
| chiP_rev |  | AAACCTGCCGCGTCGGGCATCAGAAGATGGTGAATGGTGC**TGTAGGCTGGAGCTGCTTCG** |
| Expression strain generation | | |
| chiP-pASK-IBA5_for | Cloning *chiP* from *E. coli* BW25113 into the pASK-IBA5 plasmid | ATGAATAGTTCGACAAAAATCTAGATGCGTACGTTTAGTGGC |
| chiP-pASK-IBA5_rev |  | GGTCCCCCTGCAGGTCGACCTCGAGTCAGAAGATGGTGAATGG |
| pET19b-chiP_for | Cloning *chiP* from pASK-IBA5-chiP into the pET19b plasmid | ACGTACGCATGGTATATCTCCTTC |
| pET19b-chip_rev |  | CATCTTCTGACATATGCTCGAGGATCC |
| chiP-pET19b_for |  | GAGATATACCATGCGTACGTTTAGTGGC |
| chiP-pET19b_rev |  | CGAGCATATGTCAGAAGATGGTGAATGGTG |

**Table S3.** Single channel conductance for OmpF, OmpC, ChiP, LamB in 1M KCl. pH 7±0.5

| **Channel** | Conductance KCl 1M |
| --- | --- |
| **OmpF** | 4.2 ± 0.8 nS |
| **OmpC** | 2.5 ± 0.7 nS |
| **Chip** | 0.55 ± 0.2 nS |
| **LamB** | 0.15 nS |

**Table S4.** Reversal Potential Permeability Measurements for LamB. Determined zero current potentials (V_rev_) with standard deviation and calculated permeability ratios from the mean zero current potential.

| Substrate Kanamycin  Sulphate | Substrate (mM)  Cis Trans | | Charge | V_rev_ (mV) | P_K_/P_Kan_^+4^/**P_Sulp_**_hate_^-2^ |
| --- | --- | --- | --- | --- | --- |
| K^+^ | 160 | 60 | +1 | 16±4 | P_K_/**P_Sulp_**_hate_^-2^  8:1 |
| Sulphate | 80 | 30 | -2 |  |  |
| K^+^ | 60 | 60 | +1 | -7.5±3 | P_K_/P_Kan_^+4^/P_Sulphate_^-2^  ^8:1:0^ |
| Kanamycin | 50 | 0 | +4 |  |  |
| Sulphate | 130 | 30 | -2 |  |  |
| Substrate Gentamicin  Sulphate | Substrate (mM)  Cis Trans | | Charge | V_rev_ (mV) |  |
| K^+^ | 60 | 60 | +1 | -9±5 | P_K_/P_Gen_^+4^/P_Sulphate_^-2^  ^8:1:0^ |
| Gentamicin | 50 | 0 | +4 |  |  |
| Sulphate | 130 | 30 | -2 |  |  |
| Substrate Amikacin  Sulphate | Substrate (mM)  Cis Trans | | Charge | V_rev_ (mV) |  |
| K^+^ | 60 | 60 | +1 | -6±3 | P_K_/P_Ami_^+4^/P_Sulphate_^-2^  ^8:1:0^ |
| Amikacin | 50 | 0 | +4 |  |  |
| Sulphate | 130 | 30 | -2 |  |  |

**Table S5.** Reversal potential permeability measurements ^5,6^. Determined zero current potentials (V_rev_) with standard deviation and calculated permeability ratios from the mean zero current potential. The difference in the sulfate ion concentrations is due to the titration of H_2_SO_4_ for adjusting the pH of the antibiotic solutions to 7 ± 0.5.

| Channel | (trans-side)  (mM) | | (Ground/cis-side)  (mM) | | V_rev_ (mV) | P =  P_Gen_^4+^/P_sulfate_^2-^ |
| --- | --- | --- | --- | --- | --- | --- |
|  | **Gentamicin^+^** | **SO_4_^2-^** | **Gentamicin^+^** | **SO_4_^2-^** |  |  |
| OmpF | 20 | 50 | 35 | 75 | -15 ± 3 | 1:25 |
| OmpC | 20 | 50 | 35 | 75 | -16.5 ± 3.5 | 1:30 |
| ChiP | 20 | 50 | 35 | 75 | -2 ± 1.2 | 1:3 |
|  | **Amikacin^+^** | **SO_4_^2-^** | **Amikacin^+^** | **SO_4_^2-^** | **V_rev_ (mV)** | **P_Ami_^4+^/P_sulfate_^2-^** |
| OmpF | 5 | 10 | 15 | 30 | -36 ± 3.5 | 1:10^6^ |
| OmpC | 5 | 10 | 15 | 30 | -30.3 ± 2 | 1:10^6^ |
| ChiP | 5 | 10 | 15 | 30 | -10.3 ± 3 | 1:10 |
|  | **Kanamycin^+^** | **SO_4_^2-^** | **Kanamycin^+^** | **SO_4_^2-^** | **V_rev_ (mV)** | **P_Kan_^4+^/P_sulfate_^2-^** |
| OmpF | 20 | 35 | 50 | 87.5 | −8.7 ± 2.5 | 1:14 |
| OmpC | 20 | 35 | 50 | 87.5 | −5.1 ± 0.6 | 1:4.2 |
| ChiP | 20 | 35 | 50 | 87.5 | -6.5 ± 2 | 1: 6 |

**Flux Calculation**

The single channel conductance for OmpF in Kanamycin 20 mM was 16 pS and the reversal potential for this gradient was V_rev_ = 9.6 mV which gives a Permeability ratio of 1:14. Note: Kanamycin, Amikacin & Gentamycin have the charge state charge 4^+^ and the counterion sulfate 2^-^. This makes the calculation complex. The permeability ratio should be for the flux of molecules-

1.At the reversal potential and at 20 mM we have an ion current:

I = 16 pS * 9.6 mV = 0.15 pA

Or at 10 µM Kanamycin sulfate (I/2,000)

I _total_ (1 μM) = 8 * 10^-17^ A

2.Separation into the flux of the individual ions goes via the ion current ratio:

P_Ion_ =1:14, the permeability ration is for the ion current, kanamycin has 4^+^ whereas you have 2 sulfate ions for each kanamycin

Kanamycin molecular flux is only 1/8 of the total

I_Kanamycin_ = 10^-18^A

Which gives the number of molecules by dividing the ion current by Avogadro´s number

(1 A = 1.6* 10^-19^ As)

Flux = (1 *10^-17^ A/4 (since four charges per molecule)/1.6 * 10^-19^ As = 10 molecules /sec

Note: in the table we used 10 μmol to have even numbers.
